# DksA controls the response of the Lyme disease spirochete Borrelia burgdorferi to starvation

**DOI:** 10.1101/421636

**Authors:** William K. Boyle, Ashley M. Groshong, Dan Drecktrah, Julie A. Boylan, Frank C. Gherardini, Jon S. Blevins, D. Scott Samuels, Travis J. Bourret

## Abstract

The pathogenic spirochete *Borrelia burgdorferi* senses and responds to diverse environmental challenges, including changes in nutrient availability, throughout its enzootic cycle in *Ixodes* spp. ticks and vertebrate hosts. This study examined the role of DnaK suppressor protein (DksA) in the transcriptional response of *B. burgdorferi* to starvation. Wild-type and *dksA* mutant *B. burgdorferi* strains were subjected to starvation by shifting mid-logarithmic phase cultures grown in BSK II medium to serum-free RPMI medium for 6 h under microaerobic conditions (5% CO_2_, 3% O_2_). Microarray analyses of wild-type *B. burgdorferi* revealed that genes encoding flagellar components, ribosomal proteins, and DNA replication machinery were downregulated in response to starvation. DksA mediated transcriptomic responses to starvation in *B. burgdorferi* as the *dksA*-deficient strain differentially expressed only 47 genes in response to starvation compared to the 500 genes differentially expressed in wild-type strains. Consistent with a role for DksA in the starvation response of *B. burgdorferi*, fewer CFUs were observed for *dksA* mutant after prolonged starvation in RPMI medium compared to wild-type *B. burgdorferi*. Transcriptomic analyses revealed a partial overlap between the DksA regulon and the regulon of Rel_Bbu_, the guanosine tetraphosphate and guanosine pentaphosphate [(p)ppGpp] synthetase that controls the stringent response; the DksA regulon also included many plasmid-borne genes. Additionally, the *dksA* mutant strain exhibited constitutively elevated (p)ppGpp levels compared to the wild-type strain, implying a regulatory relationship between DksA and (p)ppGpp. Together, these data indicate that DksA along with (p)ppGpp direct the stringent response to effect *B. burgdorferi* adaptation to its environment.

**IMPORTANCE:** The Lyme disease bacterium *Borrelia burgdorferi* must sense and respond to diverse environments as it cycles between its tick vectors and various vertebrate hosts. *B. burgdorferi* must withstand prolonged periods of starvation while it resides in unfed *Ixodes* ticks. In this study, the regulatory protein DksA is shown to play a pivotal role controlling the transcriptional responses of *B. burgdorferi* to starvation. The results of this study suggest that DksA gene regulatory activity impacts *B. burgdorferi* metabolism, virulence gene expression, and the ability of this bacterium to complete its natural life cycle.

## INTRODUCTION

The pathogenic spirochete *Borrelia burgdorferi* must sense and respond to its environment to complete its enzootic cycle (Samuels, 2011; Radolf et al., 2012; Caimano et al., 2016). *Ixodes* ticks acquire *B. burgdorferi* during a blood meal taken from an infected mammalian host. Thereafter, *B. burgdorferi* persist in the tick midgut through the molt. A subset of midgut-localized *B. burgdorferi* are transmitted when the next bloodmeal is acquired by the tick, which may occur up to ten months after the initial acquisition (Sonenshine, 1991; Gray et al., 2016). As *Ixodes* ticks progress through their life stages, the dynamic milieu of the midgut presents *B. burgdorferi* with multiple challenges, including variations in osmolarity, pH, temperature, nutrient availability, as well as oxidative and nitrosative stresses (Sonenshine, 1991; Bontemps-Gallo et al., 2016; Bourret et al., 2016; Caimano et al., 2016; Dulebohn et al., 2017). *B. burgdorferi* respond to changes in their environment through alterations in replication, metabolism, and outer surface protein expression (Samuels, 2011; Iyer et al., 2015; Caimano et al., 2016; Gulia-Nuss et al., 2016; Iyer and Schwartz, 2016). *B. burgdorferi* is a fastidious organism and an extreme amino acid auxotroph (Fraser et al., 1997; Gherardini et al., 2010; Groshong et al., 2017). The tick midgut following a molt and prior to a bloodmeal is a nutrient-limited and challenging growth environment for *B. burgdorferi*. Following a bloodmeal, nutrients are absorbed and sequestered away from the tick midgut. *B. burgdorferi* responds by ceasing replication and upregulating genes required to utilize available carbon sources, glycerol and chitobiose (Tilly et al., 2001; He et al., 2011; Pappas et al., 2011). Expression of genes encoding tick-associated outer membrane proteins (*ospA* and *lp6.6*) as well as nitrosative and oxidative defenses (*napA* and *uvrB*) are also required for survival and subsequent transmission (Li et al., 2007; Promnares et al., 2009; Pappas et al., 2011; Patton et al., 2013; Iyer et al., 2015; Bourret et al., 2016; Caimano et al., 2016).

The stringent response contributes to the ability of bacteria to respond to environments with limiting in nutrients. During starvation conditions, the stringent response directs resources away from cellular replication processes, including repression of ribosomal RNA synthesis, while aligning resources to maintain glycolysis and protein synthesis. The hallmark of this response is the production of the signaling molecules guanosine pentaphosphate and guanosine tetraphosphate [(p)ppGpp] from the consumption of cellular ATP and GTP or GDP (Traxler et al., 2008; Steinchen and Bange, 2016). While the production of (p)ppGpp has many consequences for the bacterium, the primary outcome is a global shift in transcription by the interaction of (p)ppGpp with RNA polymerase (Potrykus and Cashel, 2008; Hauryliuk et al., 2015). The stringent response typically results in reduced DNA replication, translation, and fatty acid synthesis, as well as increased amino acid synthesis, glycolysis, and persistence-related gene expression, which together promotes bacterial survival (Gentry et al., 1993; Paul et al., 2005; Durfee et al., 2008; Traxler et al., 2008). In *B. burgdorferi*, recent studies have shown that (p)ppGpp plays an important role in controlling expression of genes required for survival within *I. scapularis* (Bugrysheva et al., 2015; Drecktrah et al., 2015; Drecktrah et al., 2018).

*B. burgdorferi* (p)ppGpp synthetase, Rel_Bbu_, is required for the global regulatory effects of (p)ppGpp (Bugrysheva et al., 2015; Drecktrah et al., 2015). Starvation of *B. burgdorferi* in the defined culture medium RPMI induces the stringent response and a measurable increase in (p)ppGpp production (Concepcion and Nelson, 2003; Drecktrah et al., 2015). The transcriptomic response to cellular starvation provided insights into Rel_Bbu_-mediated regulation. The presence of (p)ppGpp increases the expression of genes that promote *B. burgdorferi* survival within *Ixodes* ticks, including glycerol and chitobiose utilization pathways, and *napA*. In addition, (p)ppGpp represses expression of flagellar, DNA replication, and translation-related genes, suggesting that control of these genes during starvation conditions *in vivo* is due to the stringent response. Consistent with these phenotypes, Rel_Bbu_ functions in persistence in ticks and transmission from infected nymphs to mice (Drecktrah et al., 2015).

The *B. burgdorferi* stringent response, mediated through (p)ppGpp, plays a key role in survival within *I. scapularis*; however, the role of DnaK suppressor protein (DksA) has not been investigated. DksA has emerged as an important accessory regulator of stringent responses in other bacteria (Dalebroux et al., 2010a). In *Escherichia coli*, DksA is specifically required for upregulation of amino acid biosynthesis, tRNA synthesis, and cellular utilization of alternative sigma factors (such as RpoS) that integrate the stringent response (Kvint et al., 2000; Brown et al., 2002; Paul et al., 2005; Magnusson et al., 2007; Bernardo et al., 2009; Lyzen et al., 2016). In addition, DksA holds a key regulatory role in the life cycle of several bacterial pathogens and is implicated in virulence gene expression (Dalebroux et al., 2010a; Dalebroux et al., 2010b; Pal et al., 2012; Holley et al., 2015). In enterohemorrhagic *E. coli*, DksA-dependent regulation is required for the enterocyte effacement response during intestinal colonization (Nakanishi et al., 2006; Sharma and Payne, 2006). In *Pseudomonas aeruginosa*, stringent response activation mediates colonization of surfaces by biofilm formation (Branny et al., 2001). Finally, *Salmonella enterica* requires the stringent response to respond to acidic, oxidative, and nutrient-limited environments within macrophages (Webb et al., 1999; Henard and Vazquez-Torres, 2012). In these cases, DksA works synergistically with the stringent response and is indispensable for adaptation. As seen in other bacteria, *B. burgdorferi* responds to starvation by production of (p)ppGpp, but the contribution of DksA to the regulation of the stringent response is unknown.

In this study, we expand the understanding of the *B. burgdorferi* stringent response by characterizing the role of a the DksA ortholog during adaptation to nutrient limitation. We generated a *dksA* mutant strain of *B. burgdorferi* and starved the spirochetes in RPMI medium to evaluate the role of DksA during the stringent response. Compared to BSK II, RPMI medium lacks numerous nutrients required for growth of *B. burgdorferi* including serum, oligopeptides, *N*-acetylglucosamine, along with a lower concentration of glucose. During starvation in RPMI, *B. burgdorferi* cease replication and increase synthesis of (p)ppGpp (Concepcion and Nelson, 2003; Drecktrah et al., 2015). A whole transcriptome analysis using the custom *B. burgdorferi* Affymetrix microarray chip (Iyer and Schwartz, 2016) was used to examine the response of wild-type and *dksA* mutant spirochetes to starvation. The following results indicate that starvation of *B. burgdorferi* in RPMI medium led to a DksA-dependent shift of the global transcriptome and support designating the *bb0168* gene product as a functional DksA.

## RESULTS

### Characterization of a putative DksA encoded by *bb0168*

DksA homologs are encoded in many bacterial genera, including *Borrelia*. The structure of DksA has been extensively characterized in *E. coli* (Blaby-Haas et al., 2011; Ross et al., 2016). Protein interaction studies have demonstrated that the *E. coli* DksA protein’s α-helices in the coiled-coil motif interact with the RNA polymerase secondary channel, and that the coiled coil-tip aspartic acid residues exert DksA function in the RNA polymerase core (Perederina et al., 2004; Lennon et al., 2012; Furman et al., 2013). In addition, DksA harbors a zinc-finger domain that potentially modulates its protein function (Henard et al., 2014; Crawford et al., 2016). A SWISS-model was generated for the 125 amino acid DksA protein encoded by the *B. burgdorferi bb0168* ORF using an *E. coli* RNA polymerase / DksA complex crystal structure as a template (Molodtsov et al., 2018), and visualized alongside the 151 amino acid *E. coli* DksA for comparison (**Figure 1A**). The*B. burgdorferi* DksA harbors an N-terminal 31 amino acid truncation and is nearly three kDa smaller than the *E. coli* DksA, 14.5 and 17.5 kDa respectively. The *B. burgdorferi* DksA has only 23.6% amino acid sequence identity to *E. coli* DksA; however, the SWISS-model local quality estimate indicates high similarity within the coiled-coil motif and the C-terminal region (0.6 to0.9 quality score). Moreover, an alignment of the *E. coli* and *B. burgdorferi* primary DksA amino acid sequences using the EMBOSS NEEDLE algorithm demonstrates conservation of key amino acids in DksA, including the coiled coil-tip aspartic acids in the α-helices, and the cysteines forming the zinc-finger motif (**Figure 1B**). Alignment of the amino acid sequences of DksA from various spirochetes indicates this protein is highly conserved among *Borrelia* species, suggesting a conserved function (Figure S1).

**FIGURE 1.**
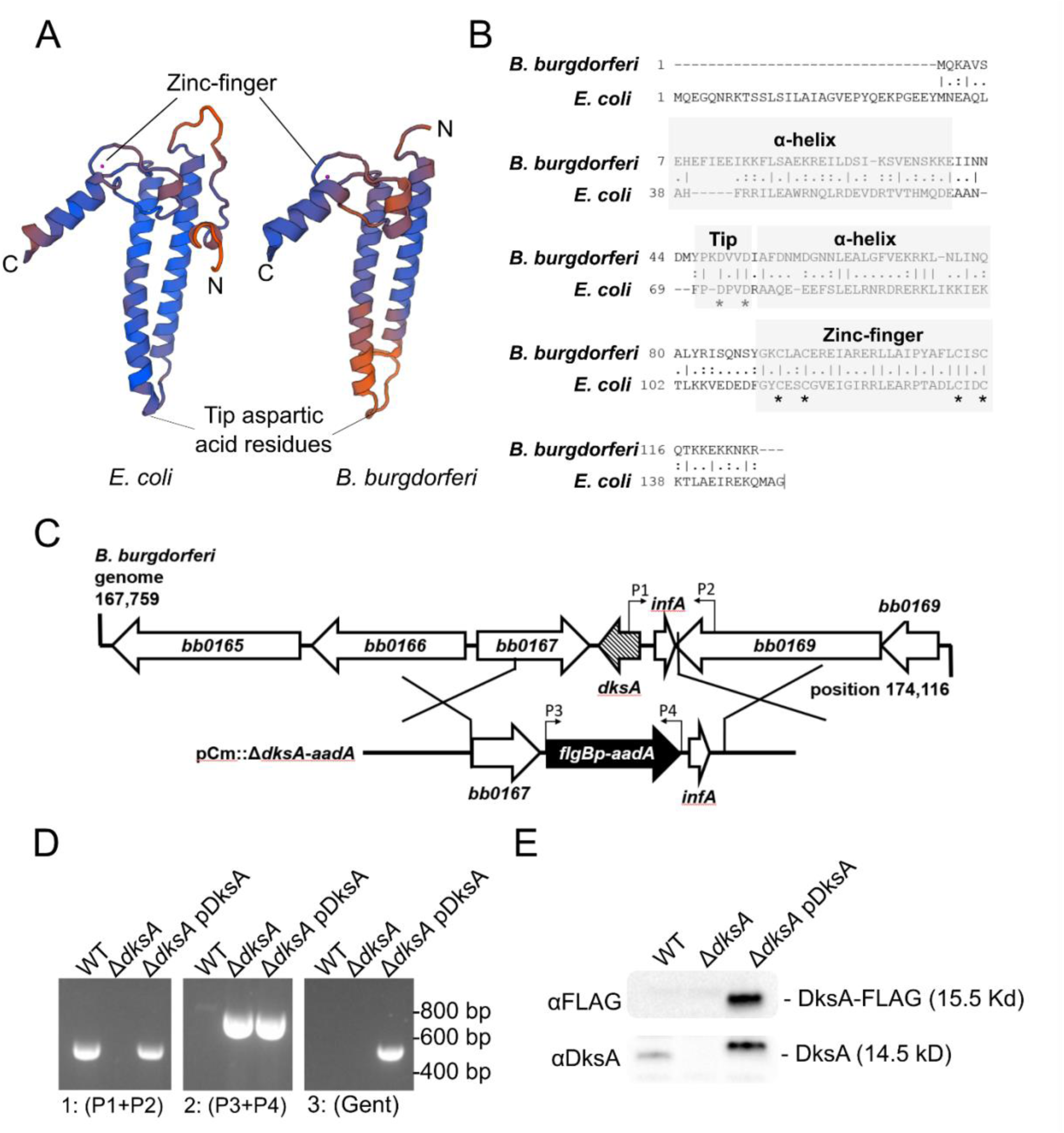
Amino acid sequence analysis and mutagenesis of conserved *B. burgdorferi bb0168*-encoded *dksA*. **(A)** SWISS-model of *E. coli* (left) and *B. burgdorferi* (right) DksA proteins illustrate predicted structural similarities. Color scale from blue (high) to orange (low) encodes Qmean score estimating model quality. Peptide N- and C-termini are indicated for each model. **(B)** Amino acid sequence alignment of *B. burgdorferi* and *E. coli* DksA protiens. The boxes indicate regions where *B. burgdorferi* DksA likely contain conserved coiled-coil α-helices and a zinc finger. The asterisks indicate key conserved aspartic acid and cysteine residues. **(C)** A schematic of the *bb0168* (*dksA*) genomic location and homologous recombination mutagenesis strategy. The open reading frame identity and direction is indicated by large arrows, and the positions of the primers used in (D) are indicated by small arrows above the genes. **(D)** Homologous recombination between the *B. burgdorferi* genome and the plasmid encoding the 600 bp segment containing *dksA*-flanking regions and the *aadA* antibiotic resistance cassette was confirmed by PCR. The δ*dksA* strain no longer possesses the *dksA* sequence as detected by PCR using the primers P1 and P2, and contains the *aadA* gene as detected with the primers P3 and P4. Additionally, the δ*dksA* strain was *trans*-complemented with the pBSV2G-based pDksA plasmid and confirmed by the presence of *dksA* detected by PCR using the primers P1 and P2, *aadA* gene detected with the primers P3 and P4, and gentamicin resistance gene (*aacC1*) with the primers *aacC1* F/R primers. **(E)** Complementation was further confirmed by western blot using antibodies targeting the FLAG (upper panel) and DksA (lower panel) epitopes.

### Generation of *B. burgdorferi* Δ*dksA* strain and *trans*-complemented Δ*dksA* pDksA strain

To study the role of *dksA* in the *B. burgdorferi* stringent response, a *dksA* mutant of *B. burgdorferi* (Δ*dksA*) was generated in the B31-A3 background. The entire *dksA* (*bb0168*) ORF was replaced by homologous recombination with a *B. burgdorferi flgB* promoter driven streptomycin resistance cassette (*flgBp*-*aadA*) used for selection (**Figure 1C**). The *dksA* mutant strain (Δ*dksA*) was complemented *in trans* with the shuttle vector pBSV2G (Elias et al., 2003) containing a *dksA* ORF fused to a sequence encoding a C-terminal FLAG epitope-tag along with 600 base-pairs of *dksA* upstream sequence (pBSV2G::*dksA*-FLAG, pDksA). The presence of the chromosomal copy of the *dksA* gene was determined by PCR (**Figure 1D**). The expression of DksA_FLAG_protein in the Δ*dksA* pDksA strain was confirmed by western blot using antibodies against FLAG and DksA epitopes (**Figure 1E**).

### Adaptation of the Δ*dksA* and Δ*rel*_Bbu_ mutants to prolonged starvation

*B. burgdorferi* wild-type and Δ*dksA* strains are morphologically similar during logarithmic growth in BSK II in microaerobic conditions (5% CO_2_ and 3% O_2_). Wild-type, Δ*dksA*, and Δ*dksA* pDksA spirochetes maintain similar maximal growth rates during logarithmic growth (**Figure 2A**). The Δ*dksA* mutant exhibited a prolonged lag phase and lower cell densities at stationary phase compared to both wild-type and Δ*dksA* pDksA strains when passaged at equivalent densities (*p*-value < 0.05). When cultures were inoculated at low densities of 1 x 10^5^ spirochetes ml^-1^, the Δ*dksA* mutants exhibited elongated morphology compared to wild-type at early time points. (Figure S2).

**FIGURE 2.**
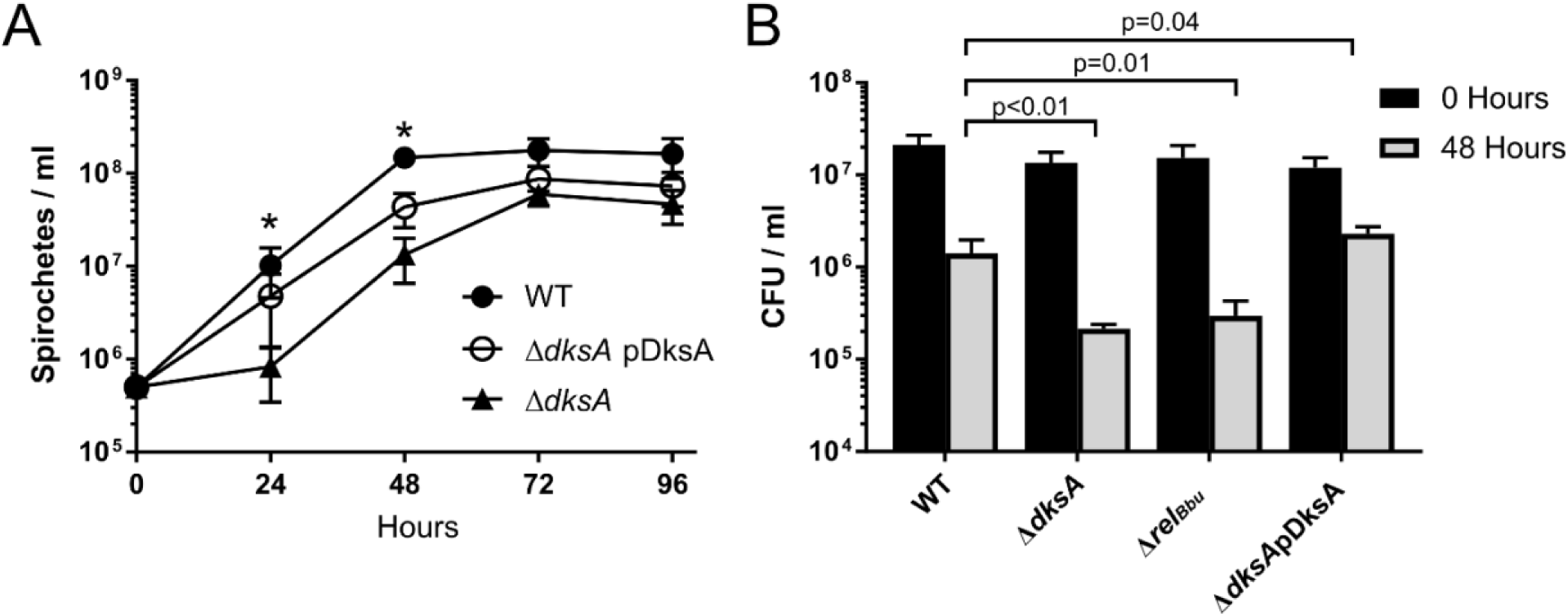
Evaluation of *B. burgdorferi* growth rate and survival during long-term starvation. **(A)** Growth of wild-type (WT), Δ*dksA*, and Δ*dksA* pDksA (passaged at 5 x 10^5^ spirochetes ml^-1^) in BSK II was assayed by enumeration at 24-h intervals. Data points represent the mean of four biological replicates. Error bars represent standard deviations and asterisks indicate *p*-values < 0.05 by one-way ANOVA. **(B)** Wild-type (WT), Δ*dksA*, Δ*rel*_Bbu_, and Δ*dksA* pDksA cultures grown to 5 x 10^7^ spirochete ml^-1^ density in BSK II were pelleted and resuspended in RPMI for 0 or 48 h prior to growth in semi-solid BSK II. Data represent the mean of three independent experiments. The *p*-values were calculated by ANOVA with a Dunnett’s multiple comparison for spirochete density following 48 h starvation in RPMI.

To determine if DksA affects survival during nutrient stress, wild-type, Δ*dksA*, Δ*rel*_Bbu_, and Δ*dksA* pDksA spirochetes were cultured to 5 x 10^7^ spirochete ml-1 and then starved in RPMI medium for 0 or 48 h and the number of colony forming units (CFUs) were assayed by plating cells in semi-solid BSK II medium. A recent study demonstrated that a *rel*_Bbu_ mutant *B. burgdorferi* (Δ*rel*_Bbu_) exhibited defect in adapting to starvation in serum-free RPMI medium (Drecktrah et al., 2015). We generated a Δ*rel*_Bbu_ strain in the B31-A3 background as described (Drecktrah et al., 2015), and consistent with previous results, Δ*rel*_Bbu_ cultures yielded significantly lower numbers of CFUs following 48 h of starvation in RPMI, compared to wild-type cultures (**Figure 2B**). Following prolonged starvation, Δ*dksA* cultures exhibited a reduction in CFUs similar to Δ*rel*_Bbu_. The Δ*dksA* pDksA restored CFUs to wild-type levels following starvation, suggesting that DksA functions in the adaptation of *B. burgdorferi* to starvation.

### Global transcriptome of the *dksA* mutant during logarithmic phase growth

To investigate DksA-dependent transcription during growth in nutrient-rich media, RNA was harvested from wild-type and Δ*dksA* mutant cultures grown to mid-logarithmic growth phase and analyzed by microarray. For these comparisons, genes were considered expressed if the hybridization signal for an ORF was significantly above background (**Figure 3A**). To evaluate differential expression, we constrained the reporting of genes to only the genes differentially expressed by two-fold (linear scale) or more, and disregarded genes when normalized, average hybridization signals were below background levels or when microarray false discovery rate (FDR)-adjusted *p*-values were 0.05 or more. The differentially regulated genes were then categorized by genomic location (chromosome or plasmid, **Figure 3B**) and function based on gene ontology (**Figure 3C**) to gain insights into DksA-dependent gene expression during logarithmic phase growth.

**FIGURE 3.**
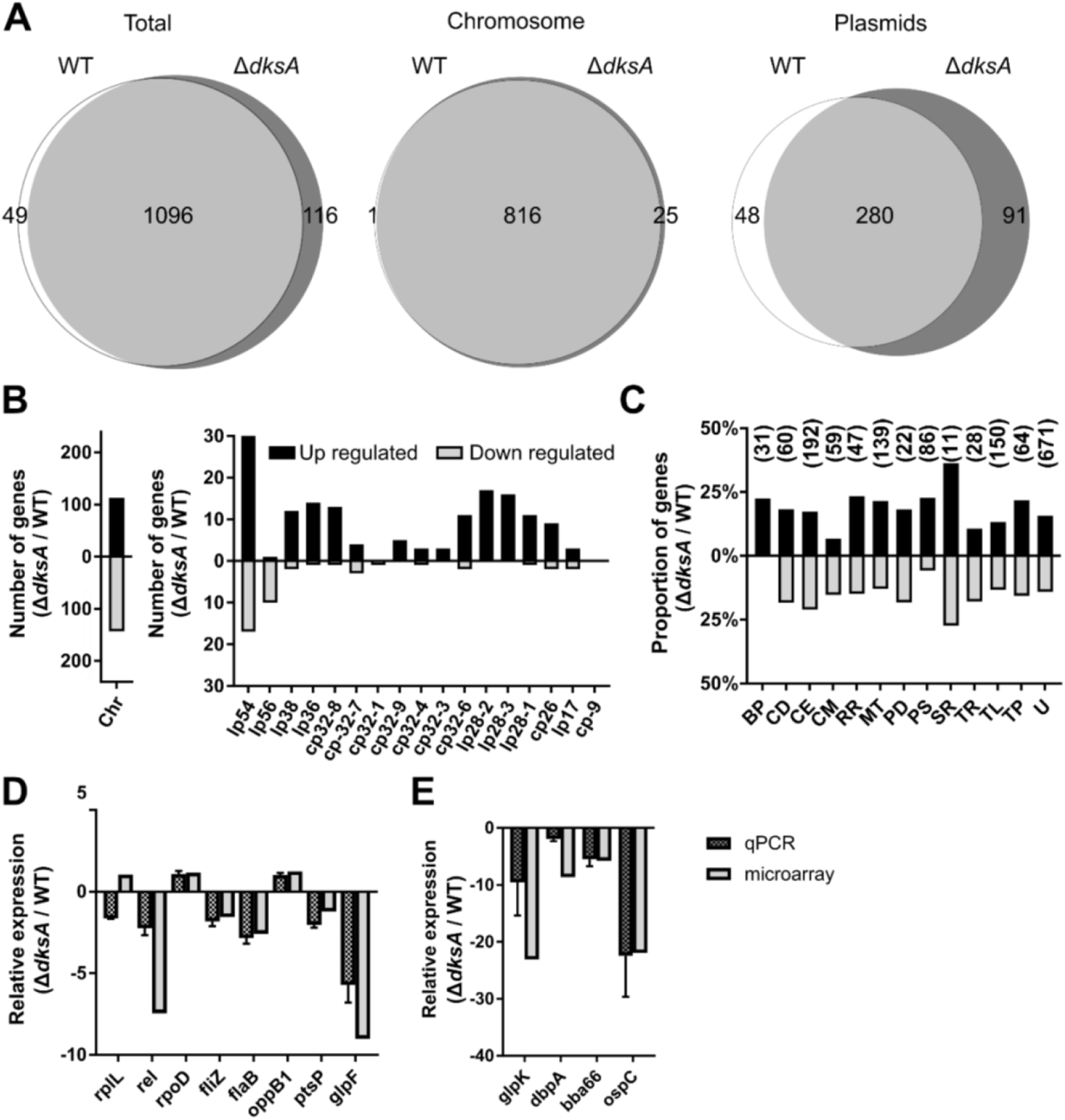
Relative RNA expression between wild-type (WT) and Δ*dksA* during logarithmic phase growth. **(A)** Venn diagrams illustrate the total number of genes expressed by WT and Δ*dksA* during mid-log phase. Expression of individual genes was determined by detection of a microarray hybridization signal above background among three biological and three intra-chip hybridization replicates (left). Genes expressed by both WT and Δ*dksA* are represented in the intersect of the two circles and the genomic location (chromosome or plasmid) is indicated (middle and right). **(B)** The number of genes upregulated (higher levels in Δ*dksA* than in WT, solid black bars) or downregulated (shaded bars) greater than two-fold and their genomic location: chromosome (Chr), linear (lp), or circular (cp) plasmids. Only comparisons with FDR-adjusted *p*-value < 0.05 are shown. **(C)** Differentially expressed genes were functionally categorized with the following abbreviations: BP, bacteriophage; CD, cell division; CE, cell envelope; CM, Chemotaxis; RR, DNA replication and repair; MT, metabolism; PD, protein degradation; PS, pseudogene; SR, stress response; TR, transcription; TL, translation; TP, transporter proteins; and U, unknown. The bars indicate percent of genes upregulated and downregulated relative to the total number of genes of each category and numbers above the bars indicate total number of genes within the respective functional group. **(D and E)** The differential regulation of select genes with high microarray signal quality or genes implicated in stringent response and infectivity were confirmed by RT-qPCR. Differential expression data by RT-qPCR and microarray are presented side by side and organized by function: ribosome (*rplL*), stringent response (*rel*_Bbu_), transcription (*rpoD, fliZ*), motility (*flaB*), transport (*bb0332, glpF*), metabolism (*ptsP, glpK*), lipoproteins (*dbpA, bba66, ospC*). Data represent the mean of four biological replicates and error bars indicate standard deviations.

During mid-logarithmic phase growth, the Δ*dksA* mutant exhibited an altered transcriptional profile compared to wild-type spirochetes, suggesting DksA is important for gene regulation during growth. The Δ*dksA* mutant expressed 1212 genes compared to 1145 genes in the wild-type (**Figure 3A**) located across the chromosome and numerous circular and linear plasmids (**Figure 3A and 3B**). The differential regulation analysis revealed that 268 genes were highly expressed in the Δ*dksA* mutant compared to the wild-type-strain (Table S1), while 186 transcripts were expressed at lower levels by the Δ*dksA* mutant (Table S2). Because both Δ*dksA* or Δ*rel*_Bbu_ mutants are susceptible to the starvation condition in RPMI, we assessed the overlap of the putative DksA and RelBbu regulon by matching genes similarly regulated by either Δ*dksA* or Δ*rel*_Bbu_. Overlap with two previous transcriptomic studies identifying genes differentially regulated in Δ*rel*_Bbu_ indicate up to 115 genes are cooperatively regulated by DksA and Rel_Bbu_ (Table S1 and S2). The genes encoding glycerol utilization proteins *glpF* and *glpK*, and oligopeptide transporters *oppA1* and *oppA2*, were similarly down regulated in the Δ*dksA* and Δ*rel*_Bbu_ mutants compared to the wild-type. The expression of genes encoding tick-associated outer membrane proteins *ospA* and *lp6.6*, and the antioxidant defense gene *napA* were also similarly regulated in the Δ*rel*_Bbu_ strain. The Δ*dksA* mutant additionally expressed genes associated with stress responses at higher levels than the wild-type strain (Table S1), including those encoding chaperones (*dnaK* and *dnaJ*), DNA repair proteins (*ligA* and *uvrB*) and numerous bacteriophage genes (*bbl01, bbn23*). In addition, the Δ*dksA* strain and Δ*rel*^Bbu^ strains both exhibit increased expression of selected genes encoding ribosomal proteins (*rpmA, rplB, rplV, rpsS, rpsC*) suggesting both (p)ppGpp and DksA are required to suppress these genes. These results are suggestive of partially overlapping regulons between Rel_Bbu_ and DksA.

To validate the microarray findings, RT-qPCR was performed comparing wild-type and Δ*dksA* spirochetes during logarithmic phase growth (**Figure 3D & E**). RT-qPCR confirmed relative expression of genes that produced high microarray signal quality (*oppB1* and *ptsP*), are implicated in the stringent response (*rel*_Bbu_, *glpF*, and *glpK*), have housekeeping functions (*rplL, rpoD*, and *flaB*), or are required for infectivity (*dbpA, bba66*, and *ospC*). Many of these genes (*rplL, rpoD, fliZ, flaB, bb0332*, and *ptsP*) are highly expressed genes with nearly 100 transcripts per 1,000 transcripts of 16S rRNA during logarithmic phase growth. Transcriptional studies have indicated that the glycerol utilization pathway is a key metabolic pathway regulated by the stringent response (Bugrysheva et al., 2015; Drecktrah et al., 2015). Two genes, *glpF* and *glpK*, encoding the glycerol transporter and kinase, respectively, were expressed at lower levels in the Δ*dksA* mutant compared to wild-type, indicating an overlap in the regulation of the glycerol utilization pathway (Table S2). Eleven of the 12 genes assayed exhibited the same direction and similar magnitude of relative expression in the RT-qPCR and microarray results. These data corroborate the findings of our microarray experiments and indicate a global effect of DksA on transcription.

### DksA mediates transcriptional responses to starvation

DksA orthologs regulate transcription in model bacteria. Therefore, we evaluated the role of DksA in the *B. burgdorferi* stringent response by comparing differences of the transcriptional responses of wild-type and Δ*dksA* strains to starvation in RPMI. For microarray analysis, RNA was harvested from cultures grown to mid-logarithmic growth phase and then following 6 h of incubation in serum-free RPMI. In wild-type spirochetes undergoing starvation, there was a dramatic reduction in the number of genes exhibiting above background microarray hybridization signals. While 1,145 genes were expressed in wild-type spirochetes during logarithmic growth in BSK II, only 587 genes were detected in wild-type spirochetes following starvation in RPMI, revealing a global reduction in transcription (**Figure 4A**). A total of 274 genes were upregulated and 226 genes were downregulated in response to starvation, indicating a restructuring of the wild-type transcriptome (Table S3), consistent with previous results obtained using differential RNA sequencing analysis (Drecktrah et al., 2015). In contrast, the Δ*dksA* mutant undergoing starvation retained the expression of the majority of genes expressed during logarithmic growth in BSK II (**Figure 4B**). Within this sizable subset of genes expressed in the Δ*dksA* mutant, only 47 genes were differentially regulated (Table S4). Thus, transcriptional remodeling of the genome during nutrient stress is to a great extent dependent on DksA.

**FIGURE 4.**
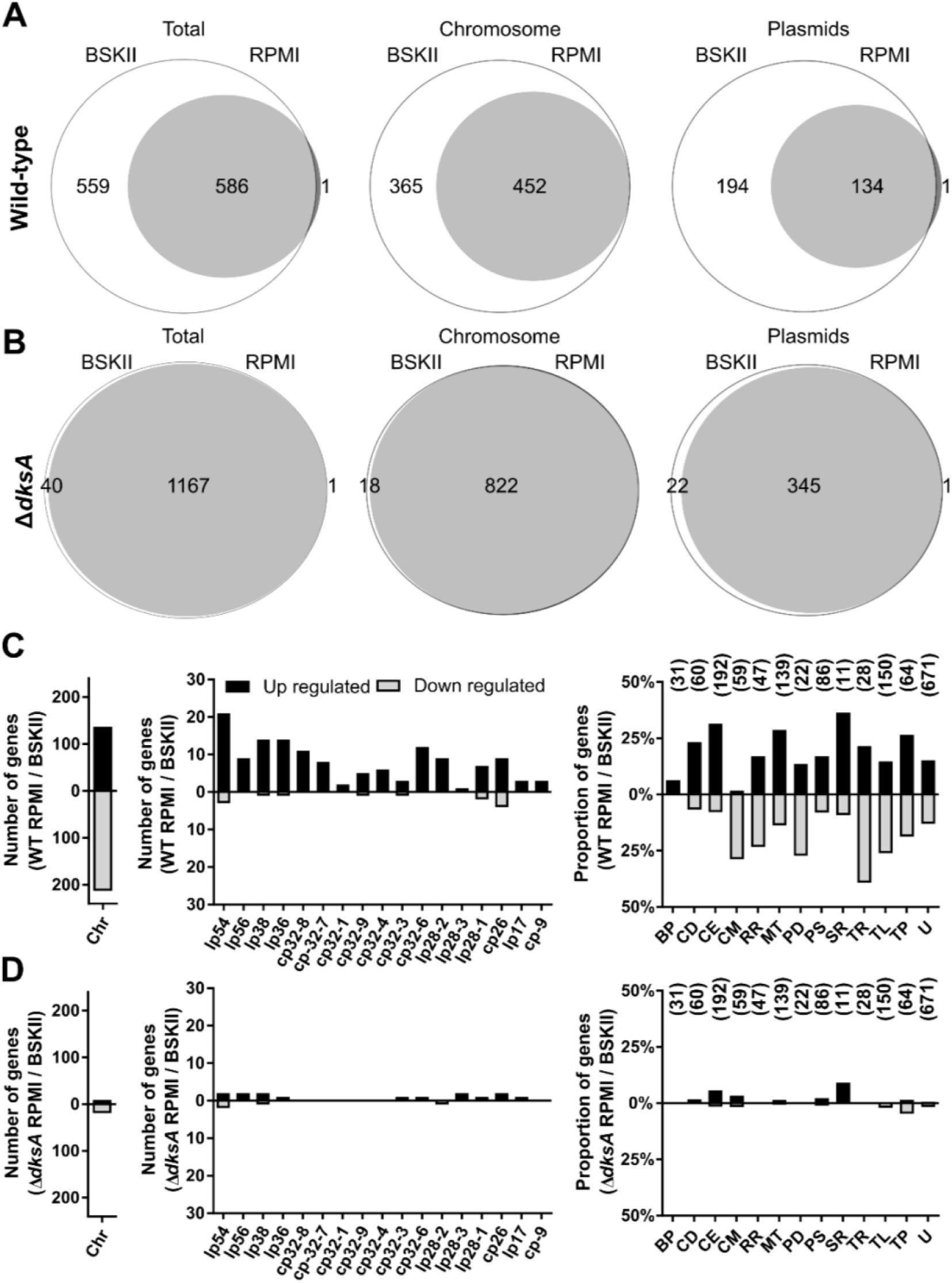
DksA mediates transcriptional responses to starvation. **(A)** The Venn diagram illustrates the number of genes expressed during mid-logarithmic (BSK II) or during starvation (RPMI) by wild-type (WT) *B. burgdorferi* or **(B)** by the Δ*dksA* strain. The data are represented as the total number of genes (left) or divided into number of chromosomal (Chr) or plasmid-encoded genes. Genes expressed exclusively during mid-log phase or during starvation are represented outside the union of the two circles, whereas the genes expressed in both arerepresented within. **(C)** The number of differentially expressed genes by cultures of WT and **(D)** Δ*dksA* strains during starvation (RPMI) as compared to mid-log phase cultures (BSK II). Bars represent the number of genes differentially expressed on the chromosome (Chr), on the various plasmids, or the percent of genes differentially expressed within the annotated functional categories relative to genes within the respective functional groups. The bars indicating proportions in the following categories: BP, bacteriophage; CD, cell division; CE, cell envelope; CM, chemotaxis; RR, DNA replication and repair; MT, metabolism; PD, protein degradation; PS, pseudogene; SR, stress response; TR, transcription; TL, translation; TP, transporter proteins; and U, unknown. Numbers above the bars indicate the total number of genes within respective functional groups. Genes were considered differentially expressed if comparisons with FDR adjusted *p*-value < 0.05 and differential expression of two-fold or more.

Genes differentially expressed by wild-type spirochetes undergoing starvation treatment were organized by gene location and functional category to characterize the transcriptional response. In wild-type spirochetes, transcriptional downregulation in response to starvation is mostly limited to chromosomally-encoded genes (**Figure 4C**). 213 of the total 226 downregulated genes were on the chromosome. Downregulated chromosomal genes are overrepresented in four functional categories: chemotaxis, DNA replication and repair, transcription (and transcriptional regulation), and translation. Among the chemotaxis genes, 13 of the 17 downregulated genes encoded flagellar components (Table S3). Genes encoding DNA replication proteins were also downregulated, including *gyrA* and *gyrB* (3.4- and 2.4-fold respectively) encoding DNA gyrase, *dnaB* (3.2-fold) encoding the replicative DNA helicase, and *dnaN* (5.2-fold) encoding the β-clamp of the DNA polymerase. The reduction in expression of DNA replication and flagellar synthesis genes is consistent with the observation that *B. burgdorferi* do not increase in CFU during starvation conditions. Additionally, we identified 39 downregulated genes encoding translation machinery including 19 genes encoding ribosomal proteins, suggesting a reduction in ribosome synthesis. A total of 17 genes in the transcription functional category were also differentially regulated during starvation. Genes encoding core transcriptional machinery were among the 11 downregulated genes, including *rpoA* and *rpoZ* (6.2-fold and 3.5-fold, respectively) encoding RNA polymerase subunits, *rpoD* (3.7-fold) encoding the housekeeping sigma factor, and *nusB* (7.6-fold) encoding the transcription anti-termination factor. Conversely, *csrA* (6.8-fold) encoding the the carbon storage regulator, *dksA* (4.4-fold), and *rpoS* (3.8-fold) encoding the alternative sigma factor were among the upregulated transcriptional regulator genes. In summary, levels of a large portion of RNA transcripts encoding crucial components of replication, transcription, and translation were decreased in wild-type spirochetes undergoing starvation. Given the functions encoded by these downregulated genes, our observations are consistent with stringent responses among bacteria. None of the genes in these four functional categories listed above were differentially regulated in the Δ*dksA* mutant, therefore the down regulation of these genes during starvation appears DksA-dependent (**Figure 4D**).

Typically, the stringent response activates the expression of genes encoding enzymes for amino acid synthesis, glycolysis, and persistence mechanisms. Consistent with the stringent response, *B. burgdorferi* undergoing starvation also upregulate genes in the functional categories of translation, metabolism, and transcription. Expression of genes that potentially increase translational efficiency were upregulated (Table S3). These genes include *infA* (4.75-fold) encoding translation initiation factor, *efp* (2.8-fold) and *tuf* (5.0-fold) encoding peptide elongation factors, and five aminoacyl-tRNA synthetases required for synthesis of Asp-tRNA^asp^,His-tRNA^His^,Ile-tRNA^Ile^,Leu-tRNA^Leu^,Val-tRNA^Val^, which recognize 33% of codons utilized by *B. burgdorferi* open reading frames (Lafay et al., 1999). The *B. burgdorferi* genome lacks many genes encoding for amino acid biosynthesis pathways and imports oligopeptides into the cell through transporters to support protein synthesis. Four oligopeptide transporter genes were upregulated; *oppA5* (6.2-fold), *oppF* (5.8-fold), *oppD* (2.5-fold), and *oppB* (2.5-fold). The transcriptome of genes involved in translation and oligopeptide transport in the Δ*dksA* mutant did not overlap that exhibited by wild-type spirochetes during starvation. Additionally, wild-type spirochetes upregulated five genes encoding enzymes involved in glycolysis during starvation: *pfk* (2.4-fold) encoding 1-phosphofructokinase, *fbaA* (2.1-fold) encoding fructose-bisphosphate aldolase, *gapdh* (5.1-fold) encoding glyceraldehyde 3-phosphate dehydrogenase, *gmpA* (5.5-fold) encoding phosphoglycerate mutase, and *eno* (5.7-fold) encoding enolase. *B. burgdorferi* lacks an electron transport chain and ferments sugars to lactate for generation of ATP. During starvation of wild-type spirochetes, no genes encoding enzymes involved in glycolysis or transporters for glucose, fructose, and chitobiose were down regulated. In contrast, the Δ*dksA* strain exhibited lower transcript levels of genes encoding key glycolysis genes enolase *eno* and pyruvate kinase *pyk* during logarithmic growth and the Δ*dksA* strain did not share the breadth of upregulation in genes encoding glycolysis enzymes in response to starvation compared to wild-type spirochetes.

### Increased expression of plasmid encoded genes in response to starvation conditions

Wild-type spirochetes undergoing starvation also differentially expressed genes carried on the numerous circular and linear plasmids (**Figure 4C**). Differentially expressed genes were largely limited to those encoding lipoproteins and hypothetical proteins, with 91% of those genes upregulated. These upregulated genes include those encoding nine OspE related proteins (*erp*) and eight multi-copy lipoproteins (*mlp*) carried on cp32s, with 3.1- to 9.8-fold and 4.8- to 13.1-fold upregulation, respectively (Table S3). In addition, *revA* (6.4-fold) and *bbk32* (2.6-fold), encoding fibronectin binding proteins, were also upregulated. Specifically, the gene product of *bbk32* regulates the classical pathway of complement and is important for infection (Lin et al., 2015; Garcia et al., 2016). The biological significance of lipoprotein regulation during starvation in RPMI medium is unknown. Overall protein expression and the level of immunogenic protein expression by wild-type and Δ*dksA* spirochetes remain relatively constant following 6 h of incubation in RPMI (Figure S3). Starvation is not thought to induce the mammalian-infection associated RpoN-RpoS cascade (Burtnick et al., 2007; Caimano et al., 2007; Samuels, 2011) and, as expected, transcription of the RpoS-regulated genes *dbpA* and *ospC* was not upregulated in response to nutrient limitation in wild-type spirochetes.

Compared to the wild-type strain, the Δ*dksA* mutant upregulated the expression of *revA, dbpA*, and *ospC* genes in response to starvation (Table S4). The Δ*dksA* mutant did not share the increased expression of *erp* or *mlp* genes with the wild-type strain during starvation. We investigated the possibility that these genes were constitutively upregulated in the Δ*dksA* mutant because the expression of many plasmid genes was higher compared to the wild-type strains during logarithmic growth (**Figure 3B**). A total of 41 plasmid-borne lipoproteins encoding genes were differentially expressed by the Δ*dksA* mutant during logarithmic growth (Figure S1 and S2). However, *revA, bbk32, erp*, and *mlp* genes had no clear pattern of constitutively higher expression by the Δ*dksA*. Moreover, we found genes encoding lipoproteins under the control of RpoS regulation, and important for *B. burgdorferi* transmission, such as *bba66* (5.8-fold), *dbpA* (8.6-fold), and *ospC* (22.0-fold) were expressed at lower levels by the Δ*dksA* mutant during logarithmic growth. Previously, the Rel_Bbu_ too appeared to control genes of the same pathway of activation: *rpoS, bosR* and *ospC* (Drecktrah et al., 2015). These results suggest DksA and the stringent response are required for the regulation of specific transmission associates lipoprotein genes.

To confirm that the disparate expression of *bba66, dbpA*, and *ospC* was DksA-dependent, the expression of these genes was compared by RT-qPCR using RNA isolated from the wild-type, Δ*dksA*, and Δ*dksA* pDksA strains during logarithmic growth and under starvation conditions (**Figure 5**). In our complemented strain, Δ*dksA* pDksA, *dksA* was over expressed, which coincided with higher levels of expression of the *bba66, dbpA*, and *ospC*. This observation supports the hypothesis that the expression of a subset of plasmid-encoded lipoproteins is either directly or indirectly dependent on DksA. Higher levels of *dksA* expression from the pDksA vector is consistent with a previous report that this plasmid vector is multi-copy (5 – 10 copies) within the cell (Tilly et al., 2006). Additionally, RT-qPCR was performed for *rpoD, fliZ*, and *ptsP* to evaluate the effects of *trans*-complementation in the Δ*dksA* pDksA strain. In the wild-type and in the Δ*dksA* pDksA strain, *rpoD, fliZ*, and *ptsP* are down regulated in response to starvation, while the Δ*dksA* mutant failed to similarly regulate these genes (Figure S4A). RT-qPCR indicated that starvation driven transcriptional regulation of these chromosomally encoded genes was restored in the Δ*dksA* pDksA strain. We also assayed for the restoration of glycerol-utilization gene expression (Figure S4B). While the Δ*dksA* mutant showed reduced levels of expression of *glpF* and *glpK* compared to the wild-type strain, the Δ*dksA* pDksA strain did not exhibit restored expression of these genes, suggesting potential intricacies in their regulation not captured by this study. These results suggest that the cellular levels of DksA have the potential to play a key regulatory role in controlling plasmid-borne gene expression in *B. burgdorferi*.

**FIGURE 5.**
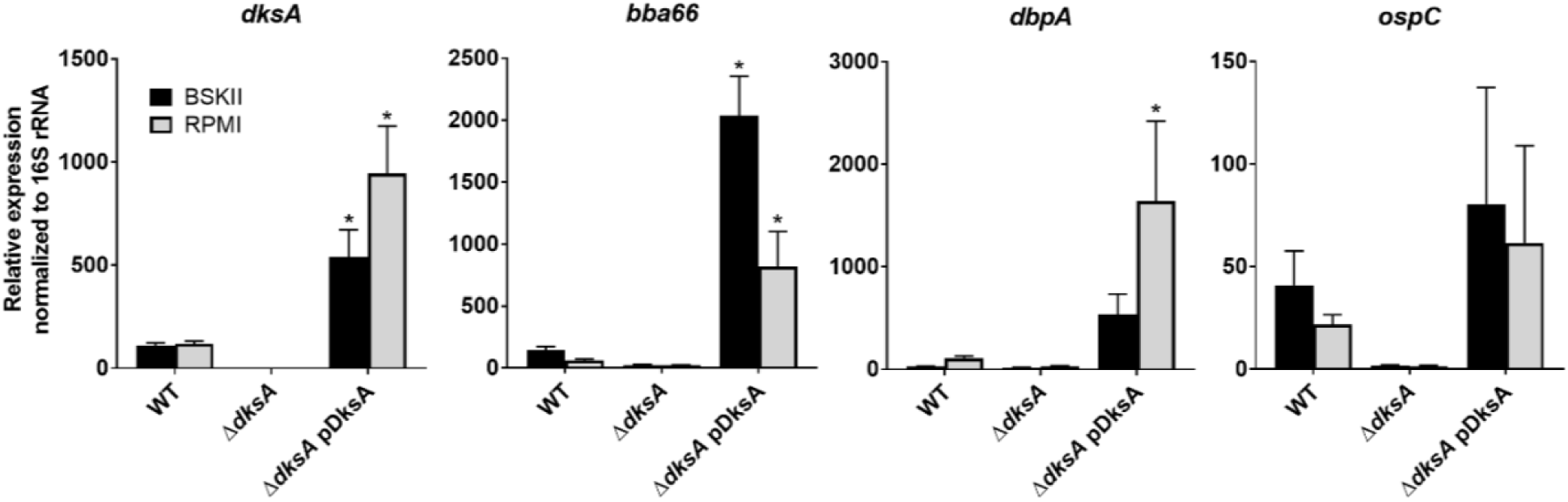
Overexpression of DksA in the Δ*dksA* pDksA strain coincides with increased expression of plasmid-encoded infectivity genes. RT-qPCR was performed on RNA extracted from wild-type (WT), Δ*dksA*, and Δ*dksA* pDksA mid-logarithmic phase cultures (black) and cultures starved in RPMI (gray). Error bars represent standard deviation calculated from four biological replicates. The Dunnett’s multiple comparison test was performed between strains for BSK II and RPMI conditions. The asterisk indicates *p*-value < 0.01 for expression level comparison between WT and Δ*dksA*, or WT and Δ*dksA* pDksA. Incubation in RPMI did not induce significant changes in expression of *dksA, bba66, dbpA*, or *ospC* for wild-type spirochetes.

### The Δ*dksA* strain overproduces (p)ppGpp

Production of (p)ppGpp and transcriptional regulation of *dksA* are intertwined in *E. coli* and (p)ppGpp also acts independently of DksA, resulting in transcriptional repression (Chandrangsu et al., 2011; Ross et al., 2016). To examine the potential interplay between (p)ppGpp by DksA, we measured the production of (p)ppGpp by thin-layer chromatography (TLC) in *B. burgdorferi* 297 wild-type, and the isogenic Δ*dksA* strain, along with the complemented Δ*dksA* pDksA strain. These 297 strains exhibit morphology, growth rate, and survival phenotypes similar to those of the respective *B. burgdorferi* B31 A3 strains (**Figure 2 and 6**). Strains were cultured to late-log (1 x 10^8^ spirochetes ml^-1^) in BSK II containing 32P-orthophosphate and nucleotides were isolated before (0 hours) or after incubation in starvation conditions (6 h) and separated by thin layer chromatography. The amount of (p)ppGpp in each strain was quantified by scanning densitometry from three independent experiments, as previously described (**Figure 7A**) (Drecktrah et al., 2015). We found the Δ*dksA* strain had significantly elevated levels of (p)ppGpp compared to the wild-type and complemented strains not only during starvation (6 Hours RPMI), but also during growth in BSK II media (0 h) (**Figure 7B**). Overproduction of (p)ppGpp inthe Δ*dksA* strain may represent a compensatory mechanism to overcome the loss of DksA. Given the 500 genes differentially regulated by wild-type spirochetes in response to starvation (Table S3), 186 of these genes were already similarly differentially expressed by the Δ*dksA* strain relative to the wild-type strain during logarithmic growth. The microarray data suggest that while the Δ*dksA* strain acts like a (p)ppGpp-deficient strain in transcription of genes encoding glycerol utilization genes, oligopeptide transporters, and ribosomal proteins, and others, the elevated levels of (p)ppGpp may play a role in the overall phenotype of the transcriptome in the Δ*dksA* strain.

**FIGURE 6.**
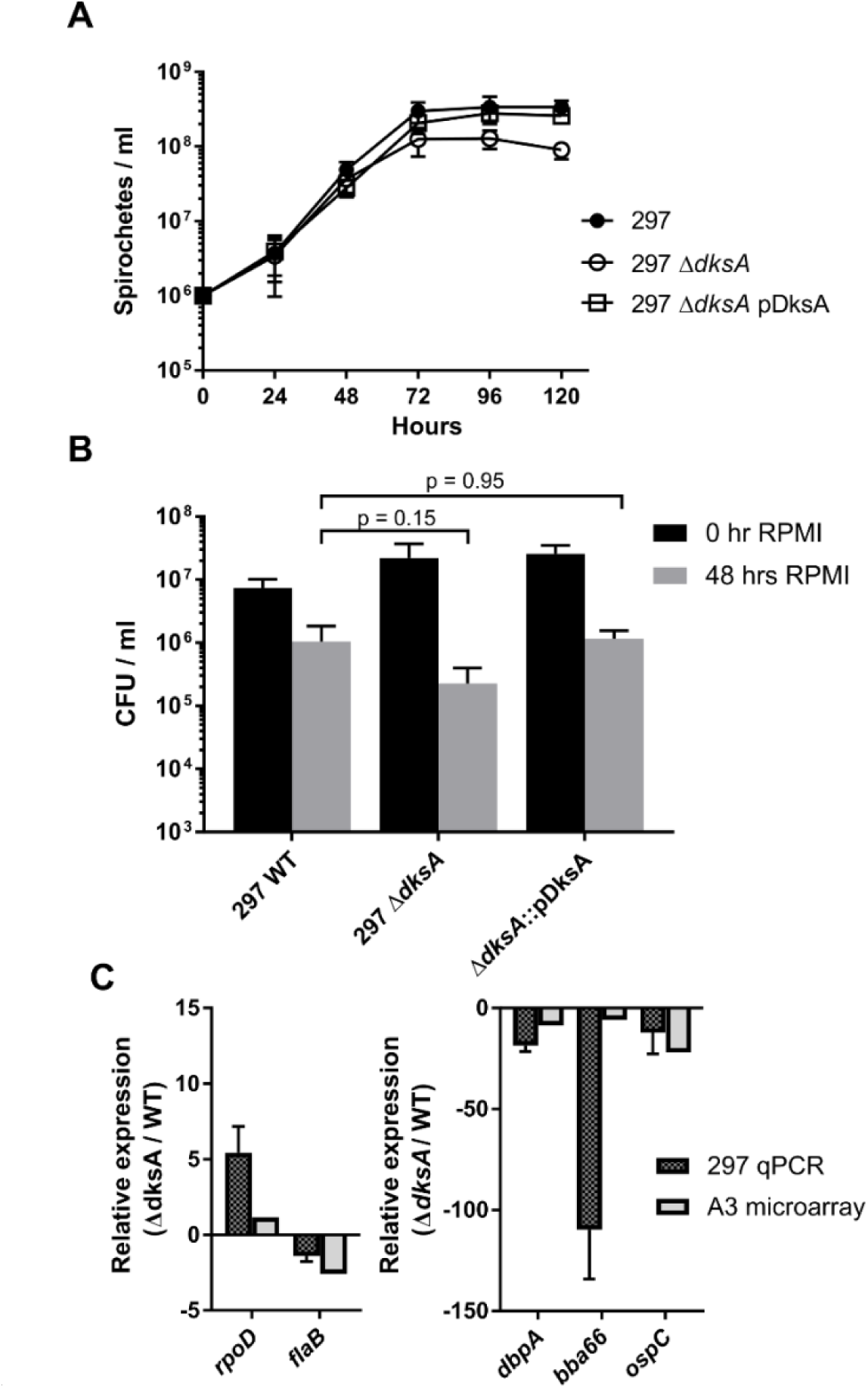
Evaluation of growth, RPMI survival, and relative RNA expression phenotypes for *B. burgdorferi* 297 wild-type (WT) and the 297 *ΔdksA* strains. **(A)** Spirochetes were enumerated by microscopy. Values represent average from two replicates and bars indicate standard deviation. **(B)** Mid-logarithmic phase cultures of 297 wild-type, Δ*dksA*, and Δ*dksA* pDksA strains grown in BSK II were pelleted and re-suspended in RPMI for 0 or 48 h before plating on semi-solid BSK II medium and CFUs were enumerated following growth. The *p*-values represent ANOVA Dunnett’s multiple comparison results of three replicate experiments. **(C)** Comparison of *dksA*-dependent gene expression in B31-A3 by microarray and 297 by RT-qPCR. Differential expression data of housekeeping genes (*rpoD* and *flaB*) and surface expressed lipoproteins (*dbpA, bba66, ospC*) are represented side by side. Relative expression values from RT-qPCR in the 297 strains represent 3 biological replicates and were normalized to 16S rRNA. Bars represent standard deviation.

**FIGURE 7.**
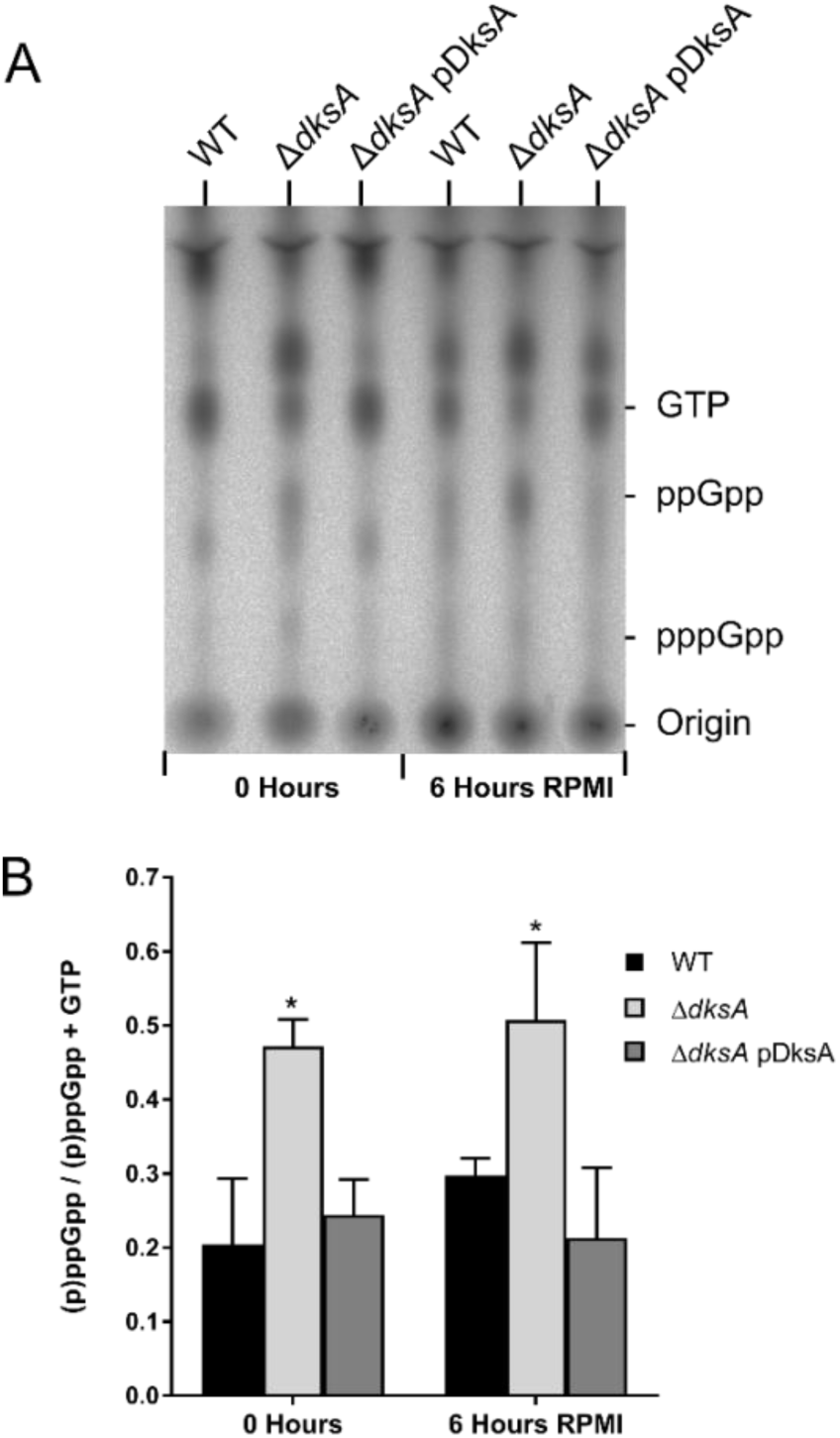
The Δ*dksA* mutant strain constitutively overproduces (p)ppGpp. **(A)** Representative TLC image for analysis of radio-labeled nucleotides from 297 wild type (WT), Δ*dksA*, and Δ*dksA* pDksA strains cultured in BSK II with _32_P-orthophosphate. Spirochetes were grown to 1 x 10^8^ spirochetes ml^-1^ (0 Hours) and starved in RPMI (6 Hours RPMI) before nucleotides were isolated and resolved by TLC. **(B)** Quantification of (p)ppGpp levels by densitometry. The values represent mean (p)ppGpp levels normalized to (p)ppGpp + GTP from three independent experiments. Error bars represent standard deviation. Asterisks indicate *p*-values < 0.05, as determined using one-way ANOVA Tukey’s post-hoc test.

## DISCUSSION

We report that the *B. burgdorferi* genome encodes a 14.5 kDa DksA protein that is involved in the transcriptional response to nutrient limitation and likely plays an additional role in controlling expression of plasmid-encoded genes. The stringent response, mediated through (p)ppGpp, is required for *B. burgdorferi* to adapt to the changes between the host and vector environments marking a shift in nutrient sources (Drecktrah et al., 2015). Therefore, we set out to characterize the role of DksA as a transcriptional regulator of the *B. burgdorferi* stringent response by simulating transition from a nutrient rich to a nutrient limited environment. Our microarray results determined that transcript levels of 500 genes changed in response to nutrient limitation (Table S3). The majority of transcriptional changes were DksA-dependent, with the expression of only 47 genes being DksA-independent under the nutrient limiting condition (Table S4). During mid-logarithmic growth, we found transcript levels of genes encoding ribosomal proteins (*rpmA, rplB, rplV, rpsS*, and *rpsC*) and stress response genes (*dnaK, dnaJ*, and *uvrB*) to be elevated in the Δ*dksA* strain, and the regulation of 41 plasmid-borne lipoprotein genes to be DksA-dependent (Table S1 and S1). The transcript levels of lipoprotein genes *bba66, dbpA*, and *ospC* were independently confirmed to be DksA-dependent in expression in 297 and A3 background strains (**Figure 5 and 6**), suggesting a pivotal role of DksA in expression of these genes. Moreover, the effects of a *dksA* deletion are likely not polar as complementation of the Δ*dksA* strain with a plasmid encoding a FLAG epitope-tagged DksA led to rescue of the Δ*dksA* phenotypes. The expression of subsets of outer surface lipoproteins in either ticks or mammals are thought to aid in the transmission and infectivity of *B. burgdorferi*. This transcriptional study provides additional evidence that the stringent response may play a role in control of outer surface lipoproteins.

Our microarray analyses suggest a partial overlap between the DksA and the (p)ppGpp regulons of *B. burgdorferi*. This is supported by the fact that the Δ*rel*_Bbu_ and Δ*dksA* mutants overlap in the lower levels of expression of oligopeptide transporters *oppA1* and *oppA2* and glycerol utilization genes *glpF* and *glpK*, while (p)ppGpp may independently regulate the glycerol utilization gene *glpD* (Bugrysheva et al., 2015; Drecktrah et al., 2015) (Table S2). The expression of the genes *ospA* and *lp6.6*, encoding tick-associated outer membrane proteins, and *napA*, an antioxidant defense gene, were reduced in Δ*dksA* mutants, suggesting the regulation of each of these requires the cooperation of DksA and (p)ppGpp. In addition, the Δ*dksA* and Δ*rel*_Bbu_ mutants both display poor adaptation to starvation since CFUs during prolonged starvation in RPMI were reduced. Wild-type spirochetes reduce transcription of replication, flagellar synthesis, and core ribosomal genes in response to starvation in RPMI medium and at the same time, upregulate genes required for peptide synthesis and glycolysis. A possible explanation for the poor adaptation to starvation by Δ*dksA* and Δ*rel*_Bbu_ mutants is the inability of the mutant strains to reduce transcription of growth- and motility-related genes to remain viable. The activity of both DksA and (p)ppGpp are likely needed for a proper response to starvation. In *E. coli*, DksA-dependent and (p)ppGpp-dependent regulation overlap to coordinate starvation-induced bacterial stringent response (Paul et al., 2004; Magnusson et al., 2007; Aberg et al., 2009; Lemke et al., 2009; Furman et al., 2015; Ross et al., 2016).

The two regulators DksA and (p)ppGpp have a close regulatory relationship in *B. burgdorferi*. Two recent transcriptional studies of *B. burgdorferi* have demonstrated that Δ*rel*_Bbu_ mutants overexpress *dksA*, suggesting that production of (p)ppGpp represses *dksA* (Bugrysheva et al., 2015; Drecktrah et al., 2015). While the role of *dksA* upregulation in Δ*rel*_Bbu_ spirochetes is unclear, we now demonstrate DksA plays a major role in transcriptional control of gene expression in *B. burgdorferi*. The transcriptomic data indicated the Δ*dksA* mutant exhibited expanded gene expression of select genes during mid-logarithmic growth and was unable to remodel the transcriptome during starvation. While the mechanism by which DksA imposes selectivity on gene transcription in *B. burgdorferi* remains to be explored, we additionally found DksA affects (p)ppGpp levels in this organism (**Figure 7**). Levels of (p)ppGpp produced in the absence of DksA were higher than levels reached by wild-type cells *in vitro* by simulating a nutrient limited condition. Moreover, Δ*dksA* spirochetes produced these levels of (p)ppGpp prior to incubation in RPMI medium, suggesting alteration of Rel_Bbu_ activity in the absence of DksA. We propose the stringent response in *B. burgdorferi* likely requires both DksA and (p)ppGpp (**Figure 8**).

**FIGURE 8.**
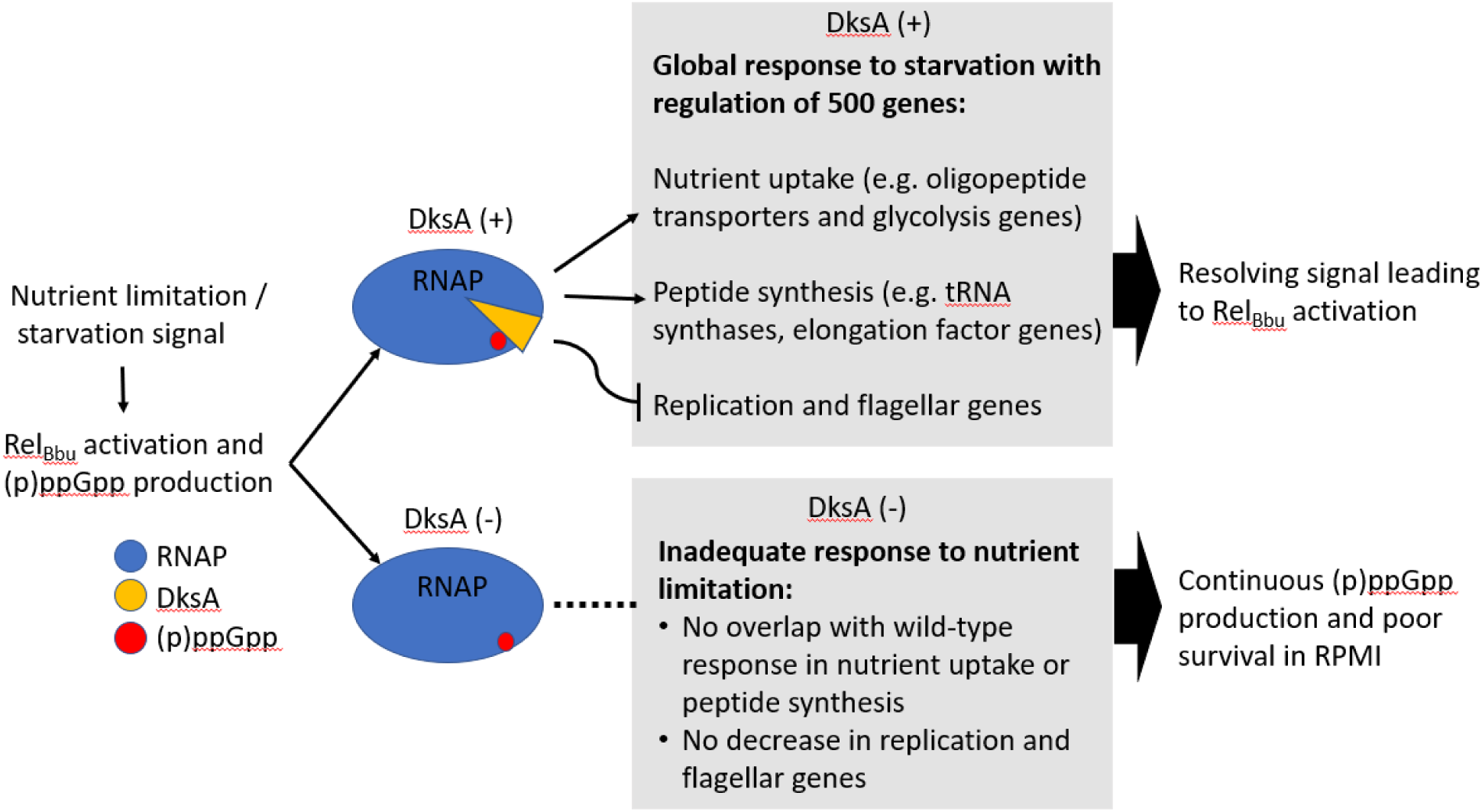
Working model of the *B. burgdorferi* stringent response. Both DksA and (p)ppGpp must interact with the RNA polymerase to exert transcriptional regulation during starvation conditions *in vitro*. In the absence of DksA, (p)ppGpp-dependent gene regulation appears largely lost, and despite the apparent overproduction of (p)ppGpp in DksA-deficient *B. burgdorferi*.

The DksA-dependent stringent response regulon potentially intersects with other regulatory mechanisms. Since (p)ppGpp is over-produced in the Δ*dksA* mutant, we cannot differentiate between the effects of (p)ppGpp from DksA-dependent regulation. (p)ppGpp is known to act independent of transcription by interacting with GTPases and riboswitches (Steinchen and Bange, 2016; Sherlock et al., 2018). Additionally, ATP and GTP homeostasis is likely altered by consumption of these nucleotide triphosphates when (p)ppGpp is produced to high levels in the Δ*dksA* mutant. All of the genes encoding xanthine/guanine permease, ribose/galactose ABC transporter, and adenine deaminases were also up regulated in the Δ*dksA* mutant (Table S1), potentially altering the flux of ATP or GTP pools. The genes encoding transmission-associated lipoproteins *cspZ, ospD, mlpD*, and *ospE* had higher expression within the Δ*dksA* mutant, the expression of which are known to be controlled by cyclic di-GMP produced by Rrp1 (Rogers et al., 2009; Caimano et al., 2015). The regulation of cyclic-di-GMP synthesis may be altered in the Δ*dksA* mutant. Additionally, transcription of the infectivity-associated lipoproteins *ospC* and *dbpA* were decreased in the Δ*dksA* mutant. The *ospC* and *dbpA* genes are regulated through a complex regulatory cascade involving Rrp2, RpoN, and RpoS (Burtnick et al., 2007; Boardman et al., 2008; Ouyang et al., 2008). As production of (p)ppGpp alters homeostasis of phosphates in bacteria (Rao et al., 1998; Hauryliuk et al., 2015), a potential point of regulatory interplay is Rrp2 (Boardman et al., 2008; Samuels, 2011). The regulation of Rrp2 phosphorylation is currently unknown, but the alteration of levels of phosphorylation in metabolic intermediates or in adenosine nucleotides may impact Rrp2 phosphorylation (Richards et al., 2015). It is possible that regulators sensitive to phosphate and nucleotide homeostasis in *B. burgdorferi* greatly contribute to the phenotype exhibited by the Δ*dksA* mutant.

In summary, we found that the *B. burgdorferi* genome encodes a DksA which contains conserved amino acid resides in the coiled-coil tip and in the zinc-finger important for DksA function in *E. coli* and *Salmonella*. Data presented here support the hypothesis that DksA is a functional transcriptional regulator in *B. burgdorferi*. We demonstrated *dksA*-dependent phenotypes in two strains of *B. burgdorferi*, B31-A3 and 297. The Δ*dksA* mutants in both B31-A3 and 297 background strains exhibit a long-term survival defect in RPMI and constitutively increased expression of housekeeping genes such as *flaB* and *rpoD*. Finally, the DksA-dependent global transcriptional changes reported here suggests DksA is fundamental for *B. burgdorferi* to adapt to environmental challenges invoking the stringent response.

## MATERIALS AND METHODS

### Bacterial strain and growth conditions

Low-passage *B. burgdorferi* B31-A3 (Elias et al., 2002) and 297 (Hughes et al., 1992) strains and their respective *dksA* and *rel*_Bbu_ mutants, and *trans*-complemented Δ*dksA* pDksA strains generated for this study were cultured in BSK II medium at pH 7.6 under microaerobic conditions (5% CO_2_, 3% O_2_) at 34 °C. BSK II media were inoculated at 1 x 10^6^ spirochetes ml^-1^ and grown to mid logarithmic phase (3 - 5 x 10^7^ spirochetes ml^-1^) density. Spirochete densities were determined by dark field microscopy, with eight microscopy fields counted per time point, and four biological replicates. Cultures from frozen stocks were passaged two times before performing assays. Construction of mutant strains is described in Supplementary Materials. The mutant strains and their plasmid profiles were determined by PCR analysis as described previously (Bunikis et al., 2011; Xiang et al., 2017) (Table S5).

### Incubation of spirochetes in RPMI

Incubation of spirochetes in RPMI and growth in semi-solid BSK II medium were performed under microaerobic conditions (5% CO_2_, 3% O_2_, 34 °C). Mid-logarithmic growth cultures were pelleted by centrifugation at 3,200 x *g* for 20 min at room temperature. The BSK II supernatant was discarded, and the pellet was resuspended in the original volume of RPMI 1640 with 2.0 mM L-glutamine (Sigma-Aldrich, St. Louis, MO, United States). The spirochetes were incubated for 6 h to compare transcription between strains or for 0 to 48 h to compare survival following long-term incubation. For quantification of viable spirochetes, *B. burgdorferi* were plated in 25 ml semi-solid BSK II medium as previously described (Samuels et al., 2018), after culture density was reduced by serial dilutions in BSK II medium.

### RNA extraction

Total RNA was extracted from 14 ml cultures at 5 x 10^7^spirochetes ml^-1^density in BSK II or RPMI. *B. burgdorferi* cells were pelleted by centrifugation at 4 °C, 3,200 x *g* for 17 min. Pellets were washed once in HN buffer (10 mM HEPES, 10 mM NaCl, pH 8.0) and then dissolved in 1 ml of RNAzol (Sigma-Aldrich, St. Louis, MO, United States) for RNA isolation according to the kit protocol. RNA integrity was confirmed by evaluation of ribosomal RNA following gel electrophoresis. The RNA was quantified by TAKE3 plate spectrophotometry in a Cytation 5 multi-mode plate reader (Biotek, Winooski, VT, USA).

### RT-qPCR analysis

cDNA synthesis was performed with approximately 1 μg of RNA with the RNA High-Capacity cDNA Reverse Transcription kit (Applied Biosystems, Foster City, CA, United States). The qPCR amplification was performed in Bullseye EvaGreen Master Mix (MIDSCI, Valley Park, MO, United States) using oligonucleotide primers specific to the gene of interest (Table S5) and detected by CFX Connect Real-Time PCR Detection System (Bio-Rad, Hercules, CA, United States). All Cq values were calculated by the CFX regression method. The Cq values of raw RNA inputs into the cDNA reaction (minus RT control) ensured that samples were DNA-free. 16S rRNA transcript levels were utilized as the reference. Typically, rRNA levels are significantly reduced during the stringent response, however, the Cq values of 16S rRNA were less responsive to varying conditions than other common reference genes such as *flaB* and *rpoD* (Figure S4). The RT-qPCR data were analyzed in Excel (Microsoft, Redmond, WA, United States) using the ΔCq method to represent transcript levels relative to 16S rRNA. Graphing and statistical comparisons were performed with Prism (GraphPad software, La Jolla, CA, United States).

### Microarray Analysis

Fragmented biotin-dUTP-labeled cDNA was prepared from purified RNA by following the Affymetrix prokaryotic target preparation protocol (Affymetrix, Santa Clara, CA, United States). The cDNA was hybridized to an Affymetrix-based Rocky Mountain Lab Custom Chip 1. Each Affymetrix chip contains three intra-chip locations for 16 antisense perfect match and mismatch probe sets against each of the 1323 ORFs of the *B. burgdorferi* strain B31 genome. One chip was used to assay for the transcriptome per biological sample. Initial quality analysis was performed on the Affymetrix Command Console version 3.1 and hybridization signals were normalized by the Affymetrix expression console version 1.1.2800 using scaling based on average cell intensity. Signal intensity principle components analysis was performed Partek Genomics Suite software v6.6 6.13.213 (Partek, Inc. St. Louis, Mo., United States) verifying that variability among biological replicates remained small compared to variability between strains and conditions. An ANOVA was performed within Partek Genomic Suite to obtain multiple test corrected *p*-values using the false discovery rate method (Benjamini et al., 2001). Fold change values and signal confidence were calculated in custom Excel templates. Importantly, our Δ*dksA* strain lacked lp-5, 21, 25, and 28-4 plasmids and the chip hybridization locations for these plasmids were excluded from the analysis.

The number of genes regulated in genomic locations or in functional categories was quantified using filters coded in RStudio (Boston, MA, United States). Affymetrix probe sets representing the gene comparisons with above background signal, ANOVA value (p < 0.05), and relative expression difference of two-fold or more were selected for representation. The number of genes that passed the criteria were totaled for each genomic segment, or alternatively, each higher or lower expressed gene was categorized by gene ontology as previously described (Bugrysheva et al., 2015; Drecktrah et al., 2015). The total gene numbers were visualized with Prism (GraphPad software, La Jolla, CA, United States).

### SDS-PAGE and immunoblot

Total cell lysates were prepared from 45 ml cultures. Spirochetes were pelleted at 4 °C, 3,200 x *g* for 17 min. Spirochetes were washed twice with HN buffer (10 mM HEPES, 10 mM NaCl, pH 8.0) and subsequently lysed in lysis buffer (4% SDS, 0.1 M Tris-HCl, pH 8.0). The lysate loading was equalized to 4 μg per sample, roughly 5 x 10^7^spirochetes, by BCA assay (Thermo Fisher Scientific, Grand Island, NY, United States). SDS-PAGE was performed on the Mini-Tetra System (Bio-Rad, Hercules, CA, United States). Proteins were detected using the EZstain system on the Gel Doc EZ imager (Bio-Rad, Hercules, CA, United States). Protein was transferred to PVDF membrane with the Transblot Turbo system (Bio-Rad, Hercules, CA, United States). The DYKDDDDK(FLAG)-tag monoclonal mouse antibody, 1 μg ml^-1^, (Thermo Fisher Scientific, Grand Island, NY, United States) was diluted 1:2000 in TBST for blotting for recombinant protein detection. Rabbit anti-DksA antibody was diluted at 1:2000 in TBST for DksA protein detection (Genscript, Piscataway, NJ, United States). Mouse serum from B31-A3 infected mice were diluted 1:200 for immunogenic protein blotting. The antibody binding was detected with the addition of HRP-conjugated secondary antibody and subsequent imaging using ECL chemiluminescence substrate (LI-COR, Lincoln, NE, United States) and the ChemiDoc Imaging system (Bio-Rad, Hercules, CA, United States).

### Measurement of (p)ppGpp

Relative quantities of (p)ppGpp were measured by TLC of radiolabeled nucleotides as previously described (Drecktrah et al., 2015). *B. burgdorferi* 297 wild-type, the isogenic Δ*dksA*, and Δ*dksA* pDksA strains were cultured to late-logarithmic growth density (1 x 10_8 _spirochetes ml_-1_) in BSK II containing 20 μCi/ml_32_P-orthophosphate (PerkinElmer, Waltham, MA, United States) in 500µl volume, pelleted by centrifugation at 9,000 x *g* for 7 min, and resuspended in RPMI. Both late-logarithmic growth density cultures (0 h) and cultures incubated in RPMI for 6 h were collected by centrifugation at 20,800 x *g* for 5 min at 4°C, cells washed once with dPBS, and cell pellet lysed with 6.5 M formic acid (Thermo Fisher Scientific, Grand Island, NY, United States). Cell debris were removed by centrifugation at 20,800 × *g* for 5 min at 4°C. The nucleotides were separated by TLC on polyethylenimine cellulose plates (EMD Millipore, Burlington, MA, United States) in 1.5 M KH_2_PO_4_, pH 3.4 buffer. After drying plates, radioactivity was detected by a 48 to 72 h exposure to an intensifying screen and screens imaged by a Fujifilm FLA-3000G Phosphorimager (Fujifilm Life Sciences, Stamford, CT, United States). Values are expressed as a ratio of ppGpp/(total ppGpp + GTP) from the densitometry of three independent experiments. Mean values from three independent experiments were analyzed using one-way ANOVA and Tukey’s post-hoc test to determine if differences were statistically significant.

## Data Availability

The microarray data have been submitted to the Gene Expression Omnibus (GEO accession: GSE119023).

## AUTHOR CONTRIBUTIONS

DD, DSS, FG, JB, WB, and TB contributed to the conception and design of the study; AG, JB, WB, and TB generated the bacterial strains required for the study; WB performed the data analysis, statistical tests, and wrote the sections of the manuscript; All authors contributed to manuscript revision, read and approved the submitted version.

## ACKNOWLEDGEMENTS

We would like to thank Rocky Mountain Laboratories Genomics unit and Dan Sturdevant for RNA expression analysis by Affymetrix Gene Chip, and Amanda Zulad, Crystal Richards, Daniel Dulebohn, and Sandy Stewart for critical review of the manuscript.

## SUPPLEMENTARY MATERIAL

The Supplementary Material for this article can be found online at:

Conflict of Interest Statement: The authors declare that the research was conducted in the absence of any commercial or financial relationships that could be construed as a potential conflict of interest.

